# Comparative analysis of convergent jellyfish eyes reveals extensive differences in expression of vision-related genes

**DOI:** 10.1101/2025.03.14.642713

**Authors:** Natasha Picciani, Cory A. Berger, Sofie Nielsen, Jacob Musser, A. Philip Oel, Marina I. Stoilova, Detlev Arendt, Anders Garm, Todd H. Oakley

**Author notes:** Natasha Picciani and Cory A. Berger contributed equally to this article.

## Abstract

Quantifying gene expression across convergent origins of traits clarifies the degree to which those traits arise from shared versus distinct genetic programs, revealing how gene re-use relates to the repeatability of evolution. Eyes are important traits that evolved in many distantly related lineages, including at least nine times within cnidarians. Here, we investigate gene expression in eye-bearing and non-visual tissues from three cnidarian species representing long-diverged lineages where eyes evolved convergently (Cubozoa, Scyphozoa, and Hydrozoa). We find gene expression in eye-bearing tissues to be mostly lineage-specific, with only a small proportion of genes having convergent expression across species. Nevertheless, all species express homologs of deeply conserved vision-related genes known from Bilateria, which likely reflects deep homology (parallel evolution across vast phylogenetic distances) of a metazoan phototransduction toolkit. A gene tree analysis of opsins—the prototypical animal photosensors—shows that convergent eyes recruited different opsin paralogs, with the potential exception of an opsin ortholog shared between scyphozoan and cubozoan eyes. Our results suggest that eyes have mostly lineage-specific patterns of gene expression, yet some key phototransduction components are repeatedly recruited across multiple independent eye origins in Medusozoa.

## 1. Introduction

Convergent evolution occurs when separate lineages evolve similar features, providing natural replicates that can clarify evolutionary processes and constraints underlying trait origins. Eyes, defined minimally as regions of photoreceptor cells adjacent to shading pigments (Arendt and Wittbrodt, 2001), have evolved many times in animals and are an excellent system to study how evolution produces sensory structures. Eyes probably evolve in the context of pre-existing light reception systems where photoreceptor cells first perform non-directional light sensing and later add components such as pigments and lenses (Nilsson, 2013; Oakley and Speiser, 2015; Picciani et al., 2021). Thus, although eyes themselves are convergent, they may recruit homologous components of an ancestral phototransduction toolkit (Oakley, 2024; Vöcking et al., 2022). This pattern is sometimes referred to as deep homology or parallel evolution (Shubin et al., 2009).

However, we do not know the extent to which eyes recruit homologous genetic components and lineage-specific novel genes because most eye origins have not been characterized at the molecular level. Eyes have evolved convergently at least nine times within medusozoan cnidarians (jellyfish), and jellyfish eyes span a range of morphologies from simple eyespots to complex lensed eyes (Berger, 1898; Conant, 1898; Miranda and Collins, 2019; Picciani et al., 2018). Examining the multiple eye origins within Medusozoa can help us gauge the extent to which convergent morphological evolution is associated with the repeated recruitment of homologous genes versus lineage-specific genetic solutions.

Eyes occur in each of the three major medusozoan classes: Cubozoa (1 origin), Scyphozoa (1-2 origins), and Hydrozoa (6 or more origins; Picciani et al., 2018). There is also a probable origin within the enigmatic Staurozoa (Miranda and Collins, 2019). Cubozoans, or box jellyfish, possess the most complex eyes among cnidarians and display visually-guided behavior (Bielecki et al., 2023); they have true image-forming eyes as well as other, simpler eye types (Nilsson et al., 2005). Cubozoan eyes are located on sensory structures called rhopalia, which contain a large part of their nervous system and perform other sensory functions (Satterlie, 2011). Scyphozoans also have rhopalia, which are probably homologous to cubozoan rhopalia (Marques and Collins, 2005). Rhopalia exist even in eyeless species and likely pre-date eye evolution. Hydrozoan jellyfish (“hydromedusae”) lack rhopalia but can possess eyes on structures called tentacle bulbs. Like rhopalia, tentacle bulbs also occur in eyeless species. Hydrozoan eyes span a range of complexity from simple pigment spots to lens-bearing eyes (Picciani et al., 2018; Weber, 1981). For most eye origins, we have no or very little information about their genetic basis.

Knowledge of eye-related genes in jellyfish is currently insufficient to compare genetic similarities among eye-bearing lineages, as only a few genes in a handful of species have been implicated in eye function and development (Ruzickova et al., 2009; Suga et al., 2010, 2008). However, it is known that both eye-bearing and eyeless cnidarians use opsin-based phototransduction mediated by cyclic nucleotide-gated (CNG) ion channels, as in Bilateria (Plachetzki et al., 2010; Vöcking et al., 2022). Genes encoding components of visual phototransduction cascades (opsins, G-proteins, and intermediary enzymes such as adenylate cyclase) and regulators of eye development (*pax*B) are expressed in box jellyfish eyes (Bielecki et al., 2014; Koyanagi et al., 2008; Kozmik et al., 2003). Knowledge of vision-related genes in Scyphozoa is limited, although *Aurelia aurita* medusae express homologs of some bilaterian eye development genes (Gold et al., 2019). Within Hydrozoa, *Cladonema radiatum* uses *pax*A and *Six*-family transcription factors in the development of its lens-eyes (Stierwald et al., 2004; Suga et al., 2010), and eye-specific opsins have also been characterized in this species (Suga et al., 2008). Various neuropeptides have been characterized in medusozoans, some of which are expressed in the rhopalia and eyes of cubozoans (Nielsen et al., 2021) and scyphozoans (Nakanishi et al., 2009). However, it remains to be seen whether these neuropeptides have specific roles in vision. Overall, genes expressed in eye tissues have not been characterized for most eye-bearing medusozoans, preventing us from identifying shared and lineage-specific genetic programs underlying eye evolution.

To address this limitation and determine the extent of genetic convergence among jellyfish eyes, we can use genome-wide approaches to gain a more comprehensive perspective on vision-related genes. Here, we compare gene expression in eye-bearing tissues from three species spanning the major clades of Medusozoa (Cubozoa, Scyphozoa, and Hydrozoa) and conduct a phylogenetic analysis of opsins in these species. We show that eye tissues (rhopalia and tentacle bulbs) from these distantly-related taxa share relatively few transcriptomic similarities, indicating that most eye-related gene expression has evolved in lineage-specific ways.

Nevertheless, we identify some homologous genes expressed convergently in the eye-bearing tissues of all three species, including members of gene families with highly-conserved roles in bilaterian vision. Our results suggest a pattern of lineage-specific innovation building on a conserved light reception toolkit, and lay the groundwork for further comparisons across the many additional eye origins within jellyfish and other animals.

## 2. Methods

### Animal culturing

We cultured polyp colonies of the hydromedusa *Sarsia tubulosa* (M. Sars, 1835) in a closed system with seawater at 18°C and 37‰ salinity as well as a 12:12 photoperiod. We obtained polyps of the scyphozoan *Aurelia aurita* sp.1 (Linnaeus, 1758) from the Santa Barbara Sea Center and kept them in an open water system with temperatures varying according to those on the coast of Santa Barbara, California. We fed polyps three-day old *Artemia* nauplii enriched with Selco (Self-Emulsifying Lipid Concentrate; Brine Shrimp Direct) every three days. *Aurelia* jellyfish were kept in a goldfish bowl with oxygenation from an air tube creating a circular flow.

### RNA extraction for tissue-specific library construction and sequencing

We starved jellyfish for at least 24 hours prior to dissections in order to avoid brine shrimp contamination. We dissected tentacles, tentacle bulbs/rhopalia and manubria (Fig. 1A-C) using UV-sterilized tools further treated with RNase AWAY™ Surface Decontaminant (Thermo Fisher Scientific). We transferred each dissected tissue directly into either the RNeasy Mini Kit lysis buffer or chilled TRIzol Reagent (Invitrogen) (see Table S1 for details on tissue samples and their RNA extraction methods). We proceeded according to the kit manufacturer’s protocol for samples extracted with the RNeasy Mini Kit. For those kept in TRIzol, we proceeded with a liquid nitrogen freezing step followed by maceration with a mini pestle and RNA extraction using chloroform. We sent RNA samples to Novogene Corporation (Sacramento, CA) for quality control tests (quantitation and RNA integrity checks using Nanodrop, Agarose Gel electrophoresis, Agilent 2100), library preparation and paired-end Illumina Hiseq PE150 sequencing of 150 bp reads. We sequenced a total of 15 paired-end RNA-seq libraries from four tissue types from *Sarsia* and *Aurelia,* with at least two biological replicates per tissue per species [*Sarsia*, tentacles: 2 replicates (32 and 40 tentacles from 8 and 10 jellyfish), tentacle bulbs: 2 replicates (32 and 40 tentacle bulbs from 8 and 10 jellyfish), manubrium: 2 replicates (8 and 10 manubria from 8 and 10 jellyfish); *Aurelia*, rhopalia: 3 replicates (31, 67, 67 rhopalia), and manubrium: 6 replicates (6, 3, 3, 9, 6, manubria; one replicate with number of manubria not recorded)]. For comparative analyses, we used tissue-specific paired-end RNA-seq data from tentacles (SRR8101526), rhopalia (SRR8101523), manubrium (SRR8101518), and whole medusa without tentacles, rhopalia, or manubrium (SRR8101525) of the box jellyfish *Tripedalia cystophora* Conant, 1897, publicly available in the NCBI SRA database (Bioproject PRJNA498176).

**Figure 1.**
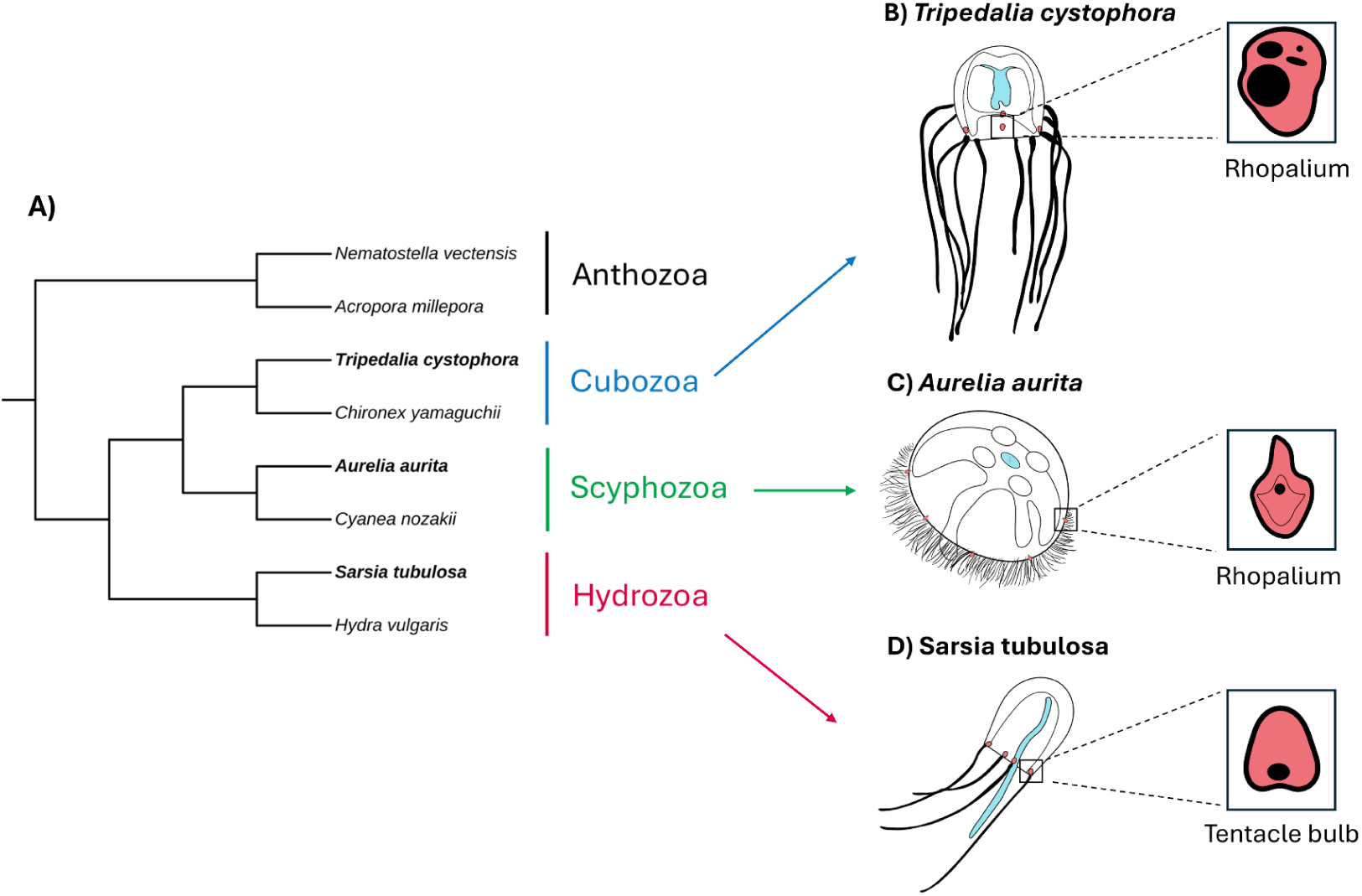
Diagrams of study species and tissue types. A: Cladogram of study species (bold) and other species used for OrthoFinder analysis. B-D: Sketches showing body plans of the study species highlighting the locations of eye-bearing tissues (rhopalia and tentacle bulbs), manubria, and tentacles. Eye-bearing tissues are colored red and shown in the inserts, with eyes represented as black ovals. Manubria (mouth and stomach) are colored cyan, and tentacles are drawn with solid black lines. Drawings are not to scale.

### Assembly of a low-redundancy reference transcriptome for Sarsia

We extracted RNA from whole body *S. tubulosa* jellyfish using the Qiagen Total RNA Kit and stranded mRNA 150 bp paired-end sequencing using Illumina NovaSeq500. We evaluated the quality of raw reads using FastQC v0.11.9, and used Trinity v2.8.5 for assembling a combined set of one whole body jellyfish library and three tissue-specific libraries (one library of each tissue; tentacles, tentacle bulbs and manubrium). According to quality assessment via BUSCO v3.1 (Seppey et al., 2019), this original assembly possessed 96.3% Metazoa Complete BUSCOs, with a high proportion of duplicated BUSCOs (53.5%) among complete BUSCOs. These high levels of duplicated genes in the reference assembly led to many multi-mapped reads in preliminary analyses, sometimes comprising almost half of the total number of mapped reads.

Multimapped reads can lead to overestimation of sequenced molecules, and, as such, are often discarded prior to differential expression analyses at the gene level [see (Deschamps-Francoeur et al., 2020)]. We lowered transcriptome redundancy by keeping only one transcript (the transcript coding for the longest protein) per ‘gene’ (’gene’ as defined by Trinity gene ids) following (Liang et al., 2019). This non-redundant transcriptome reference had 93.4% Metazoa Complete BUSCOs and 1.9% complete and duplicate BUSCOs. We used the same strategy to build a reference assembly for *Tripedalia*, which also lacks a genome, based on its original assembly available at the NCBI TSA database [GHAQ00000000.1; (Khalturin et al., 2019)]. By doing that, we lowered the number of sequences in our initial *Sarsia* and *Tripedalia* reference assemblies from 251,436 and 154,192 sequences, respectively, to 29,710 and 20,979. This step increased the downstream amount of uniquely mapped reads from ∼40-45% to ∼75-85%.

### Read mapping and counting

We used STAR v2.7.5b (Dobin et al., 2013) to build an index and perform read mappings of tissue-specific libraries (see Table S2 for STAR alignment summary statistics). We counted reads mapped to genes using featureCounts v2.0.1 (Liao et al., 2014) with an annotation GTF file containing only exon entries generated with a custom python script. In order to map reads from *Aurelia*, we used the genome sequence from *Aurelia aurita* var. Pacific Ocean published by (Khalturin et al., 2019) as a reference.

### Differential gene expression analysis

We used estimated fragment counts from featureCounts to perform pairwise contrasts to identify genes differentially expressed (DE) among tissues using the R package DESeq2 (Love et al., 2014). For each pairwise contrast, the independent filtering optimizes removal of low expressed genes in order to maximize the number of rejections (Benjamini Hochberg adjusted p-values lower than α=0.05) over the mean of normalized counts. Each pairwise contrast was performed using Wald statistics as implemented in DESeq2, and genes were considered DE if the adjusted p-value was less than 0.05. Because the *T. cystophora* samples have no biological replicates, we cannot confidently estimate differential expression. However, for exploratory purposes, we used NOISeq (Tarazona et al., 2015), which simulates technical replicates, to compare expression among *T. cystophora* libraries. For NOIseq, we considered genes DE if they had a q-value greater than 0.99.

### Orthology assignment and inter-species comparisons

We used OrthoFinder v2.5.5 (Emms and Kelly, 2019) to infer homology relationships. In addition to our three focal species, we provided OrthoFinder with five additional assemblies: two outgroups (*Nematostella vectensis* jaNemVect1.1 and *Acropora millepora* Amil_v2.1), and three medusozoans belonging to the same classes as each of our three focal species: Hydrozoa (*Hydra vulgaris* PRJNA497966), Scyphozoa (*Cyanea nozakii* SRR2089748), and Cubozoa (*Chironex yamaguchii* PRJNA498204). We did this because the inclusion of additional taxa to break up long branches can improve orthology inference in the species of interest (Emms and Kelly, 2019). For the public RNA-seq data, we assembled high-quality transcriptomes using the following methods: we trimmed raw reads with Trimmomatic v0.39 (Bolger et al., 2014) using default parameters, performed kmer correction with RCorrector v1.0.6 (Song & Florea, 2015), and removed ribosomal RNA with SortMeRNA v4.3 (Kopylova et al., 2012). We assembled transcriptomes using Trinity v2.15.1 (Grabherr et al., 2011) with default parameters and clustered transcripts at 99% identity using CD-HIT-EST v4.7 (Li & Godzik, 2006). We predicted peptides using TransDecoder v5.7.1 (https://github.com/TransDecoder/TransDecoder), retaining peptides with hits to UniRef90 (downloaded March 8, 2024) searched by DIAMOND BLASTP v2.1.8 in “sensitive” mode (evalue = 1e-5; Buchfink et al., 2015) or hits to PFAM (downloaded March 8, 2024) as searched by hmmscan (-E 1e-10; http://hmmer.org). Finally, we used CD-HIT to cluster 100% overlapping peptides (-c 1.0) and assessed transcriptome completeness with BUSCO v5.6.1 against the Metazoa odb10 database, downloaded January 11, 2024 (Simão et al., 2015).

The assemblies had BUSCO scores of 94.2% (*C. nozakii*), 94.8% (*H. vulgaris*), and 95.5% (*C. yamaguchii*). We analyzed phylogenetic hierarchical orthogroups at the node corresponding to the common ancestor of the three focal species (Fig. 1D). We tested the significance of overlaps in DE orthogroups with the SuperExactTest package (https://github.com/mw201608/SuperExactTest/), with the background consisting of all orthogroups with at least one gene present in all three species.

To quantify gene expression similarity, we calculated expression values within each species as TPM10K, which is similar to transcripts-per-million (TPM) except that it also accounts for differences in assembly size among species (Munro et al., 2022). For principal components analysis (PCA) and hierarchical clustering, TPM10K values were log-transformed with a pseudocount of 0.001 and quantile normalized (results were similar with or without quantile normalization). PCAs were performed with the ‘prcomp’ R function and hierarchical clustering was performed on Pearson correlations with the complete linkage method using the ‘heatmap.2’ R function. Samples were clustered based on the set of 2682 single-copy orthologs shared among the three species. As we initially observed strong clustering by species, we also applied Z-score normalization to the expression values and re-performed clustering analyses, enabling us to compare relative changes in expression. We used the “betadisper” function in the ‘vegan’ R package to test for differences in group dispersion between eye tissues and manubria in PCA space (first 19 PC’s).

### GO enrichment analysis

We performed functional annotation of all three transcriptomes using EggNOG-mapper v2.1.12 (Cantalapiedra et al., 2021). We tested whether Gene Ontology (GO) terms under the “Biological Process” category were significantly enriched among upregulated genes with the Fisher’s exact test with a GO processing algorithm (“weight01”) that takes into account the hierarchical nature of gene ontology terms using the R package topGO (Alexa et al., 2006), and a p-value cutoff of 0.01. We tested the significance of GO term overlaps between species with the SuperExactTest package, with the background consisting of all GO terms found among the tested genes of all species.

### Phylogenetic analysis of opsin sequences

We retrieved opsins from *Sarsia*, *Aurelia* and *Tripedalia* references using a customized python version of PIA [Phylogenetically Informed Annotation; (Speiser et al., 2014)] with the opsin tree from Picciani et al. (2018).

## 3. Results

### Eye-bearing tissues do not cluster together across species

We generated RNA-seq tissue libraries for *Aurelia aurita* (manubria and rhopalia) and *Sarsia tubulosa* (manubria, tentacles, and tentacle bulbs), and re-analyzed published data for *Tripedalia cystophora* (manubria, tentacles, rhopalia, and whole medusa without manubria, tentacles, and rhopalia; Bioproject PRJNA498176). Although the *T. cystophora* tissue libraries did not have replicates, they are included as a further point of comparison. In principal components analyses (PCA) within each species, samples clearly separated by tissue type (Fig. 2A; Fig. S1), and eye-bearing tissues were distinct from adjacent tentacles in *S. tubulosa* and *T. cystophora*.

**Figure 2.**
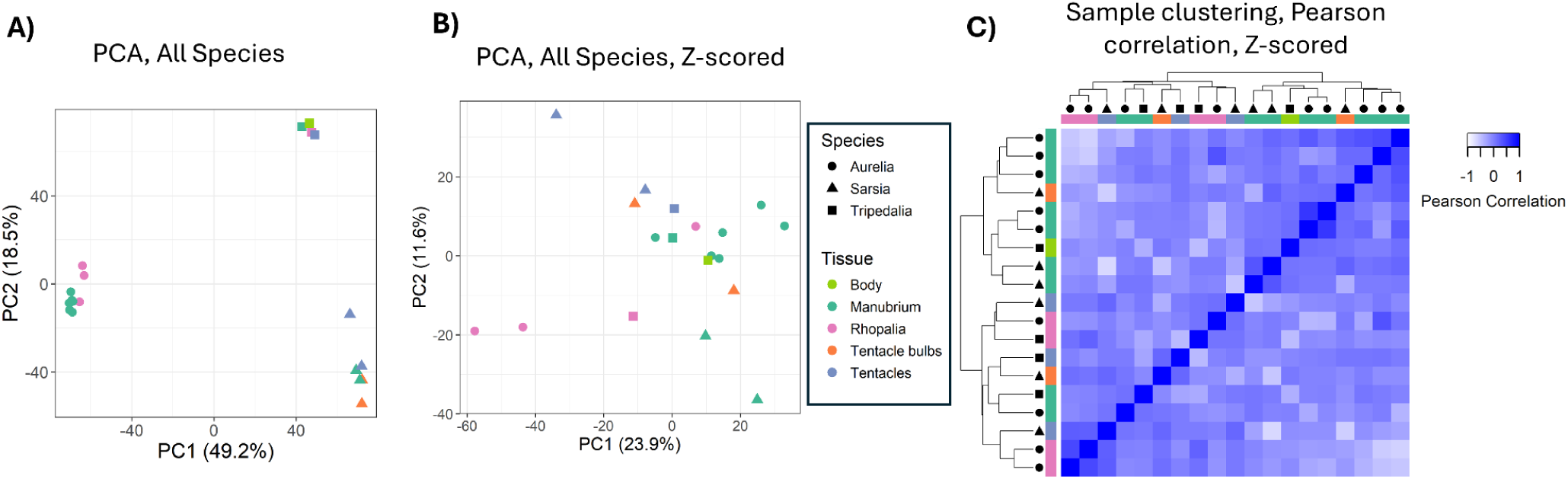
Gene expression signatures and clustering of tissue samples. A: Principal component analysis (PCA) of tissue samples across species, calculated from expression of 2682 single-copy orthologs with quantile normalization. Expression data were expressed as log2 TPM10K with a pseudocount of 0.001 and analyzed with the ‘prcomp’ R function. Numbers in parentheses indicate the proportion of variance attributable to each principal component. B: Same as in A, except quantile-normalized expression values were Z-scored. C: Sample clustering based on Pearson correlations of samples (quantile-normalized, Z-scored log2 TPM10k of single-copy orthologs).

Cluster dendrograms on the top and left display hierarchical clustering of tissue samples using the complete linkage method, and shapes indicate species.

Considering all species together, samples clustered strongly by species rather than tissue (Fig. 2A), based on the expression of 2682 single-copy orthologs. In a PCA, the first principal component (49% of the variance) separated *A. aurita* from the other two species, whereas the second principal component (19% of the variance) separated *S. tubulosa* and *T. cystophora* (Fig. 2A). Clustering of gene expression by species rather than tissue has been observed in various invertebrates, including other cnidarians (Breschi et al., 2016; Mantica et al., 2024; Munro et al., 2022). PCAs and hierarchical clustering of Z-scored data no longer clustered by species and had partial clustering by tissue type (Fig. 2B, C). Eye-bearing tissues did not generally cluster together, except for the *T. cystophora* rhopalia sample and one of the *A. aurita* rhopalia (Fig. 2C). In contrast, *S. tubulosa* tentacle bulbs clustered with manubria samples and the *T. cystophora* tentacle sample (and did not cluster with one another).

For the Z-scored data, eye tissue expression was more divergent across species than manubria expression. In other words, eye tissues (rhopalia and tentacle bulbs) had greater variance in PCA space than manubria samples (betadisper, p=0.027). Thus, gene expression was less similar among likely non-homologous tissues (rhopalia and tentacle bulbs) than tissues with likely homology (manubria), despite the convergent evolution of eyes within rhopalia and tentacle bulbs.

### Functional similarities are driven by mostly non-homologous gene sets

To characterize gene expression signatures of eye-bearing tissues (rhopalia and tentacle bulbs), we inferred differentially expressed genes (DEGs) between eye-bearing tissues, manubria, and tentacles within species (Wald test, p_adj_ <0.05). DEGs and their annotations are in Table S3.

For cross-species comparisons, we focused on differential expression between eye-bearing tissues and manubria. We identified 250 GO terms in the “biological process” category enriched among upregulated eye-tissue DEGs in *A. aurita*, 208 in *S. tubulosa*, and 80 in *T. cystophora*.

DEGs from each pairwise comparison are shown in Table 1 and all enriched GO terms are in Table S4. Broadly speaking, GO terms enriched for the eye tissue vs. manubria comparison were related to cell signalling, nervous system function, and sensory processing in all species. Each species had several GO terms specifically related to light perception, such as “detection of visible light” (all 3 species), “phototransduction” (*A. aurita* and *T. cystophora*), and “visual perception” (*A. aurita* and *S. tubulosa*).

**Table 1.** Number of up- and downregulated DEGs in pairwise contrasts between tissue types in each jellyfish species (Wald test, p_adj_ <0.05).

|  | *Sarsia* |  | *Aurelia* |  | *Tripedalia* * |  |
| --- | --- | --- | --- | --- | --- | --- |
|  | up | down | up | down | up | down |
| **Rhop/TBulbs vs Tent** | 1614 | 1815 | n/a | n/a | 359 | 516 |
| **Rhop/TBulbs vs Manu** | 2149 | 1403 | 4339 | 2798 | 494 | 444 |
| **Tent vs Manu** | 3192 | 2659 | n/a | n/a | 281 | 360 |
| **Total unique DEGs** | 7589 |  | 7137 |  | 1516 |  |
| **%DEGs out of total genes** | ~26% |  | ~24% |  | ~7% |  |

Rhop: rhopalia, TBulbs: tentacle bulbs, Tent: tentacles, Manu: manubrium, n/a= not available
\**T. cystophora* libraries did not have replicates and were analyzed with NOIseq; these results should be viewed as exploratory. NOISeq is not deterministic and depends on the starting seed.

Overlaps of GO terms between species were significantly greater than expected by chance alone (SuperExact test, p<<0.01), for all pairwise comparisons and the three-way intersection. Eleven GO terms were shared between all three species (Table 2). Eleven additional GO term was shared between *S. tubulosa* and *T. cystophora* (including “neurotransmitter transport” and “negative regulation of calcium ion transport”), and 15 between *A. aurita* and *T. cystophora* including “phototransduction”, “cGMP-mediated signaling”, “cAMP-mediated signaling”, and “response to light intensity”). An additional 32 terms were shared between *A. aurita* and *S. tubulosa*, including “regulation of circadian sleep/wake cycle, non-REM sleep”, “neuronal signal transduction”, “calcium ion transmembrane transport”, and “visual perception”. The higher number of similar GO terms in *A. aurita* and *S. tubulosa* probably reflects the fact that the *T. cystophora* analysis is exploratory and lacks statistical power.

**Table 2.**
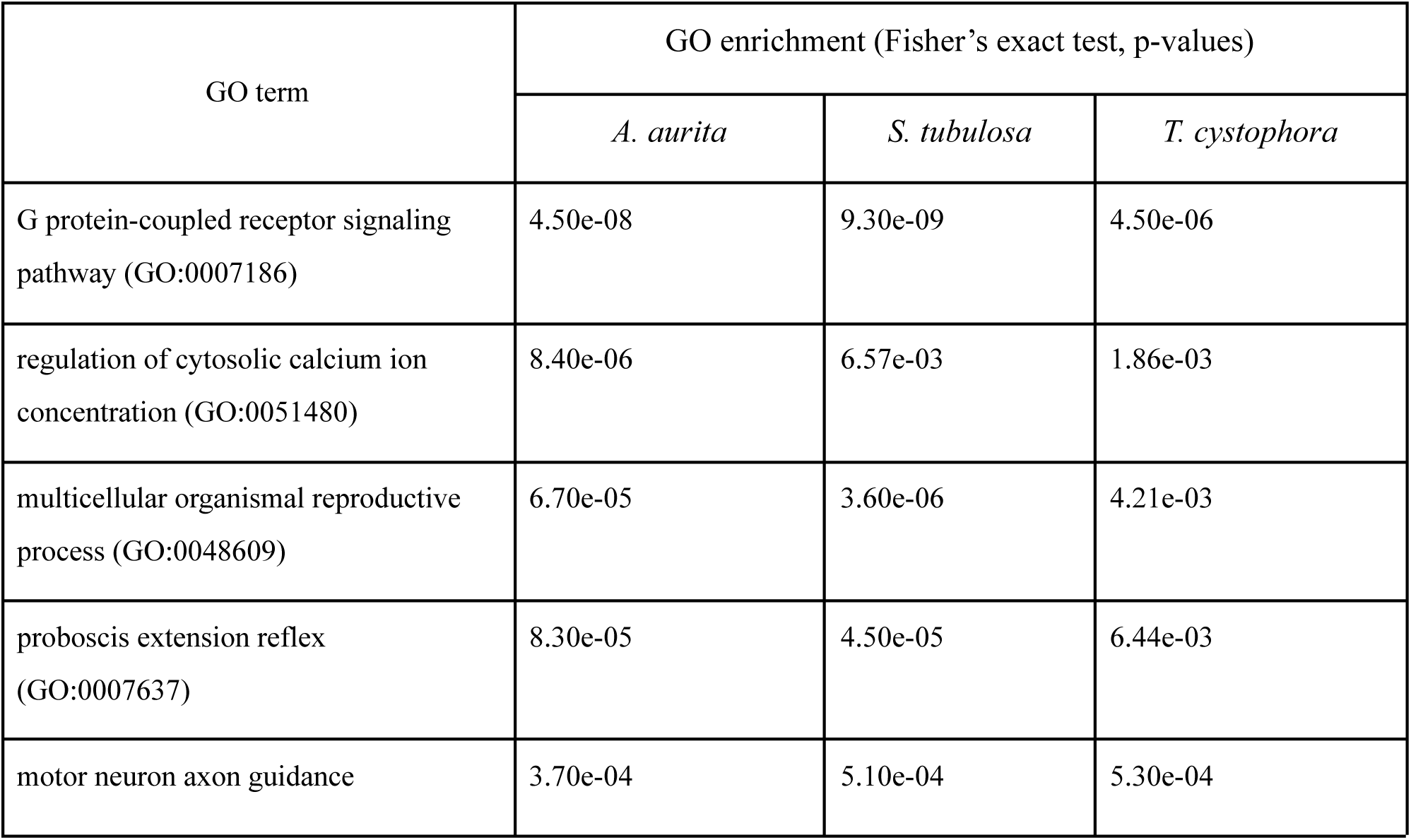
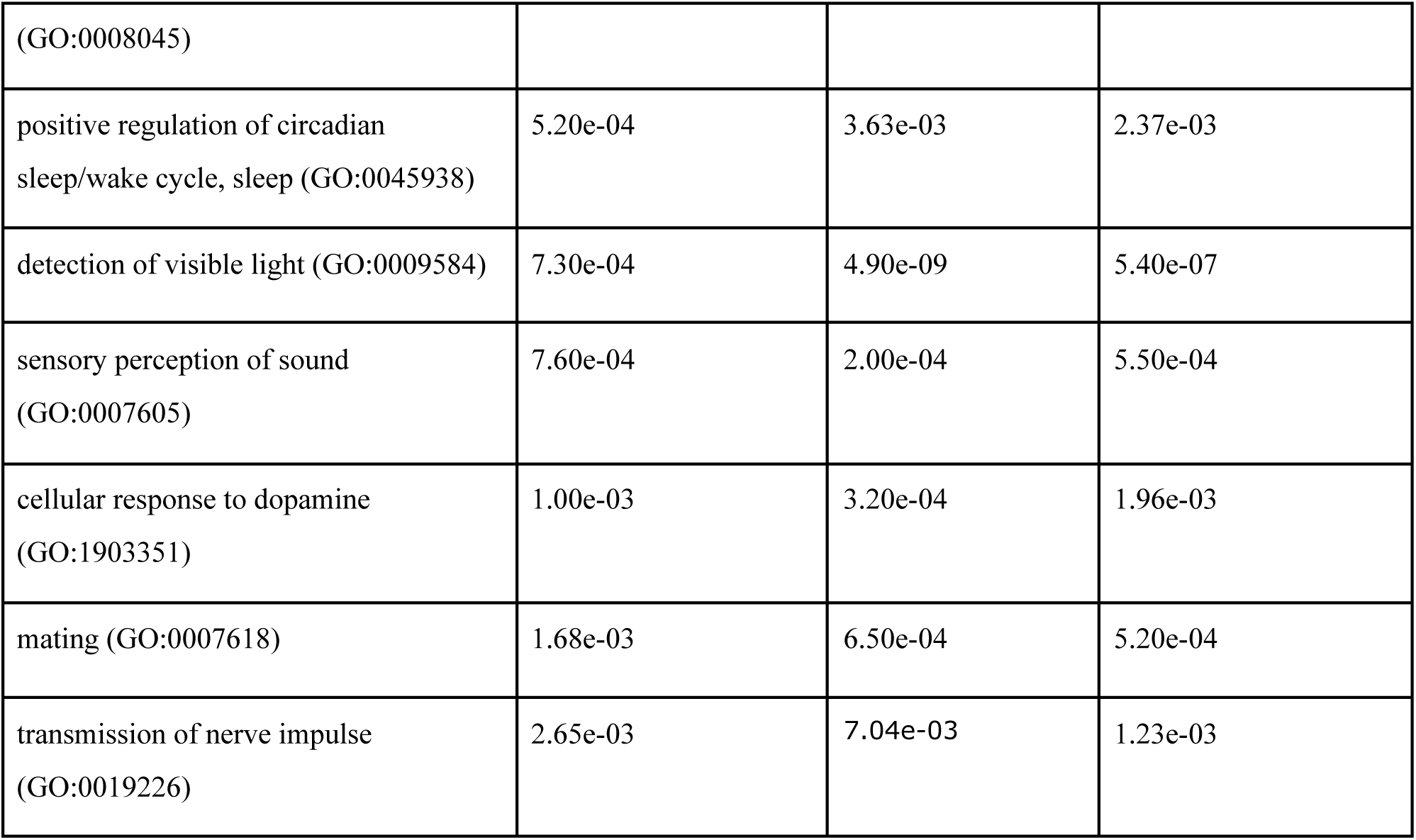
Overlapping GO terms from eye tissues shared between all three species.

As always, GO terms in non-model systems should not be interpreted too narrowly, but these results highlight the analogous functions of eye-bearing tissues in these taxa, consistent with the roles of rhopalia and tentacle bulbs as centers of sensory processing. Furthermore, the enrichment of GO terms related to vision and light perception means that these jellyfish express members of gene families with known roles in bilaterian vision. The enrichment of GPCR signaling is likely also a signature of opsin-based phototransduction, a type of GPCR signaling pathway.

Genes driving enrichment of shared GO categories across species were mostly non-homologous. Across all GO terms shared between multiple species, an average of only 24.2% of eye-tissue DEGs annotated with a given term were homologous—i.e. they belonged to an orthogroup also containing a DEG in another species (13.9% in *A. aurita*, 24.7% in *S. tubulosa*, and 39.4% in *T. cystophora*). Thus, functional similarities between eye-bearing tissues are primarily driven by lineage-specific expression of genes that are either not homologous, or that belong to large gene families split into multiple orthogroups. Nonetheless, there is a core set of homologous genes with convergent expression that contributes to a proportion of shared GO enrichment across species.

### Convergent gene expression highlights well-known vision-related genes in multiple eye lineages

Some genes with canonical roles in phototransduction were upregulated in the eye tissues of multiple species, notably including opsins, voltage-gated ion channels and other components of phototransduction pathways, and transcription factors with roles in bliaterian eye development.

Out of 2682 single-copy orthologs, 19 were upregulated in the eye-bearing tissues of all three species compared to manubria. These genes included a calmodulin, an FEZ family zinc finger protein, a cytochrome P450, a dystroglycan, a Rho GTPase-activating protein, a potassium channel, and an unannotated homeobox protein. This is a general set of genes with roles in gene regulation and signal transduction. Some of these genes have roles in nervous systems of other metazoans (e.g., calmodulins and dystroglycans; Jahncke and Wright, 2023; Solà et al., 2001), and homeobox genes function in eye development (along with development of many other structures) in both invertebrates and vertebrates.

Nine additional orthologs were upregulated in the rhopalia of *A. aurita* and *T. cystophora*, including a putative neuropeptide receptor, a voltage-dependent calcium channel subunit, and a protein kinase. A further 27 genes were upregulated in *A. aurita* rhopalia and *S. tubulosa* tentacle bulbs, including several calcium, sodium, and potassium channels; a hippocalcin-like protein (which in vertebrates is a neuron-specific calcium-binding protein); a calsyntenin; and additional transcription factors. Finally, six more orthologs were shared between *S. tubulosa* and *T. cystophora*, including an adenylate cyclase, a Kelch-like protein, and a (non-opsin) GPCR.

Extending our analysis from single-copy genes to members of the same orthogroup (which can be orthologs or paralogs), we found 25 multi-copy orthogroups containing at least one gene upregulated in eye tissues of all three species. Additionally, 49 multi-copy orthogroups were shared between *A. aurita* and *T. cystophora*, 82 between *A. aurita* and *S. tubulosa*, and 49 between *S. tubulosa* and *T. cystophora*. The numbers of overlapping DE orthogroups were no greater than expected by chance given the number of shared orthogroups between the three species (SuperExact test, p=0.16). In *A. aurita, S. tubulosa*, and *T. cystophora*, 6%, 11%, and 37% of DEG’s belonged to an orthogroup with at least one DE gene in another species, respectively (including single-copy orthologs).

Among the orthogroups shared in all three species, there were a few genes with potential roles in phototransduction or eye development: several opsins, a CNG ion channel, and a six3/6 (optix) homeobox gene. There were also numerous other transcription factors of various families, several GPCRs with similarity to neurotransmitter or hormone receptors, and a 5’-cyclic phosphodiesterase. Multi-copy genes shared between *A. aurita* and *T. cystophora* included additional opsins, a nuclear receptor, guanylate cyclases, and aquaporins, while those shared between *S. tubulosa* and *T. cystophora* included a Sox transcription factor, a guanylate cyclase, and a lens fiber membrane intrinsic protein (Lmip). Multi-copy genes shared between *A. aurita* and *S. tubulosa* included additional calcium and potassium channels, calcium-binding proteins, GPCRs, SOX transcription factors, a nuclear receptor similar to a vertebrate photoreceptor-specific NR (NR2E), guanylate cyclases, and Lhx homeobox proteins.

Finally, we examined potential vision-related genes that were only DE in a single species. In *A. aurita*, this includes additional ion channels, two cGMP-dependent phosphodiesterases, a G-protein, a Regulator of G-protein signaling gene (RGS), and a Wnt gene. *Tripedalia cystophora* rhopalia upregulated three crystallins (2 of which were previously characterized as J1 lens eye crystallins in this species; Piatigorsky et al., 1993), a gene similar to a retinal homeobox protein Rax, another CNG channel, and a homolog of a bone-morphogenic protein (BMP). Lastly, *S. tubulosa* tentacle bulbs upregulated another possible Lmip gene.

### Neuropeptides are broadly expressed across tissue types

Six neuropeptide candidates have been annotated from the transcriptome of *T. cystophora*, (Nielsen et al 2019), three of which (RF-, RA- and VWamide) are expressed in various neuronal subpopulations both within and outside the rhopalia (Nielsen et al. 2021). We investigated whether the remaining neuropeptides in *T. cystophora*, along with neuropeptides in *S. tubulosa* and *A. aurita*, are also expressed in eye-bearing tissues based on existing annotations in these species (Koch and Grimmelikhuijzen, 2019; Nielsen et al., 2019). For *A. aurita*, Koch & Grimmelikhuijzen (2019) used a different reference transcriptome, so we BLASTed those sequences against our assembly. We identified 3/6 *A. aurita* neuropeptides among our protein-coding genes. All three investigated *A. aurita* neuropeptides were greatly upregulated in rhopalia—although they were also highly expressed in manubria. In *T. cystophora*, the six neuropeptides were highly expressed in all tissues and had by far the highest expression in rhopalia; three (LWamide, RAamide, and FRamide) were DE. Of the nine *S. tubulosa* neuropeptides investigated, two (LWamide and WLKGRFamide) were upregulated in tentacle bulbs compared to manubria, but both also had high expression in tentacles. RPRamide and FFPRamide were upregulated in manubria.

### Opsins expressed in eye-bearing tissues are orthologs of known visual opsins

Because opsins are the quintessential light sensors in animal eyes, we investigated the expression levels and evolution of this gene family. Using phylogenetically informed annotation (PIA; Speiser et al., 2014) and manual inspection of BLAST hits, we identified 6, 28, and 20 candidate opsins in *A. aurita*, *S. tubulosa*, and *T. cystophora*, respectively. In the latter two species, several of these were partial-length sequences lacking the lysine residue (K296) to which the photosensitive chromophore attaches (Hargrave et al., 1983). These short sequences were excluded from further analysis, resulting in final sets of 25 and 13 opsins in *S. tubulosa* and *T. cystophora*, respectively. Consistent with previous studies, we found that opsin genes from each species all belong to the cnidops clade (Fig. 3A), the only type of opsins known from medusozoans (Picciani et al., 2018).

**Figure 3.**
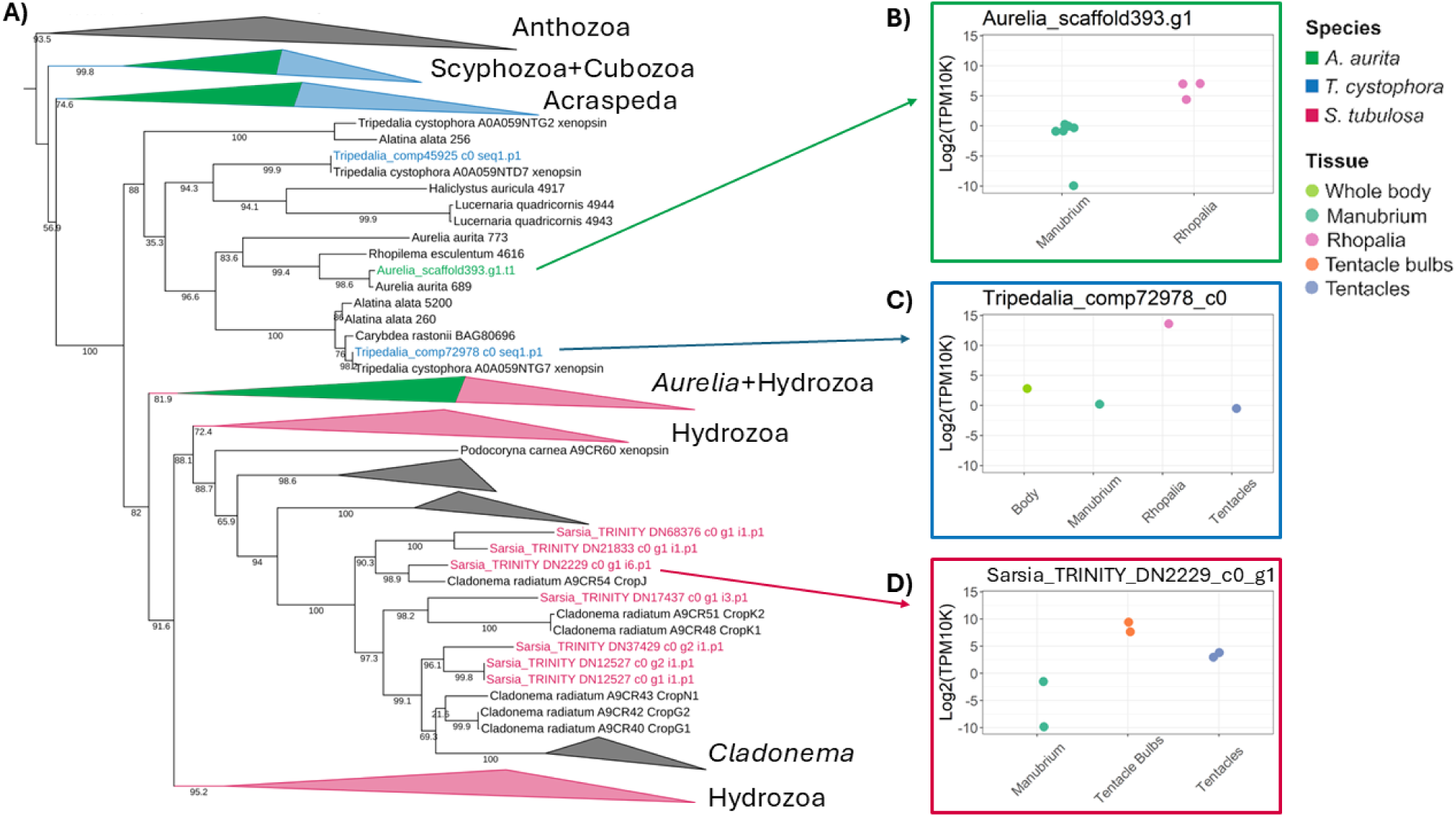
Cnidops gene tree (A) and expression of selected opsin genes (B-D). Maximum likelihood tree inferred from a sequence alignment of anthozoan and medusozoan “cnidops” sequences under the JTT+F+R8 model in IQ-TREE. Ultrafast bootstrap support values calculated from 1000 replicates shown above branches. Clades are collapsed for visualization, with groups containing opsins from *A. aurita*, *T. cystophora*, and *S. tubulosa* colored green, blue, and pink, respectively. Full tree is in the Supplemental Material

Medusozoan visual opsins have been characterized in the lens eyes of *T. cystophora* (comp72978_c0; referred to as lens-eye opsin (LEO) in Bielecki et al. (2014)) and in the eyes of the hydromedusa *Cladonema radiatum* (Suga et al., 2008). The lens-eye opsin had by far the highest expression in the *T. cystophora* dataset, with expression 4 orders of magnitude higher in rhopalia than manubria. In total, 10/13 *T. cystophora* opsins were differentially expressed, all of which had dramatically higher expression in rhopalia than manubria; most of these had low or negligible expression in manubria and tentacles, including even the three opsins that did not meet the statistical criteria for differential expression. These results were consistent with localization of opsins by antibody staining (Garm et al., 2022): the lens-eye opsin and another eye-specific opsin, the “slit-eye opsin” (comp45925_c0), were both only expressed in rhopalia. In *A. aurita*, 4/6 investigated opsins were DE, all of which had higher expression in the rhopalia and low-to-negligible expression in manubria. In the gene tree, *A. aurita* opsins fell into three groups: nested in a clade with scyphozoan and staurozoan sequences; sister to another scyphozoan opsin in a clade with cubozoan opsins (including the lens-eye opsin of *Tripedalia*); and in a clade with hydrozoan sequences. The opsin orthologous to the lens-eye opsin of *T. cystophora* had the second-highest rhopalia expression of the *A. aurita* opsins (Fig. 3B).

In *S. tubulosa*, patterns of opsin expression across tissues were more variable. Four opsins had high tentacle bulb-specific expression (upregulated in tentacle bulbs compared to both manubria and tentacles), while four were upregulated in both tentacle bulbs and tentacles. Two opsins had the highest expression in manubria, and one had the highest expression in tentacles. Thus, in *S. tubulosa*, not all opsins are associated with eye-bearing tissues, whereas all *A. aurita* and *T. cystophora* opsins were primarily expressed in rhopalia.

Strikingly, the *S. tubulosa* opsin with the strongest tentacle bulb-specific expression is sister to the CropJ eye opsin from *C. radiatum* (Fig. 3A, D). Two other *S. tubulosa* opsins are also closely related to CropJ: one of these was upregulated exclusively in tentacle bulbs, and another was upregulated in both tentacle bulbs and tentacles. CropK1 and CropK2 are other opsins expressed specifically in the ocelli of *C. radiatum*; the *S. tubulosa* ortholog of CropK1/K2 also had tentacle bulb-specific expression. Thus, there is a clade of opsins in *S. tubulosa* and *C. radiatum* that have strong eye-specific expression in *C. radiatum* and tentacle bulb-specific expression in *S. tubulosa*. *Cladonema* and *Sarsia* belong to separate families, but ancestral state reconstructions suggest that their eyes are homologous (Picciani et al., 2018). The observation that opsin orthologs have strong tentacle-bulb/eye-specific expression in both species provides further evidence for the homology of eyes in these two hydrozoan families. A clade of *C. radiatum* opsins with expression in other body parts is sister to the eye-related opsins and does not have orthologs in *S. tubulosa*. A separate clade of *S. tubulosa* opsins form a clade with *C. radiatum* opsins that have tentacle-specific expression (Suga et al., 2008); these were all either non-DE or had high expression in both tentacles and tentacle bulbs. This clade of opsins thus does not appear to have eye-specific expression but is expressed more broadly in tentacles and tentacle bulbs in both species. The remaining *S. tubulosa* opsins, some of which were highly expressed in the manubria or were not DE, were closely related to other hydrozoan sequences but did not have orthologs in *C. radiatum*.

## 4. Discussion

Separate eye origins in moon jellyfish (*Aurelia aurita*), box jellyfish (*Tripedalia cystophora*), and hydromedusae (*Sarsia tubulosa*) are associated with expression of some familiar animal eye gene families. This suggests that deeply-conserved components of an ancestral phototransduction toolkit play functional roles in convergently-evolved eyes. However, the vast majority of gene expression in eye-bearing tissues is lineage-specific. Across species, gene expression was more dissimilar among eye-bearing tissues than among manubria, which is thought to be an homologous tissue (Hejnol and Martindale, 2008; Kraus et al., 2015). These results suggest some level of convergence coupled with rampant differences in vision genes upregulated in eye-bearing tissues across species, consistent with their phylogenetically independent assemblies.

Our results highlight the roles of cubozoan and scyphozoan rhopalia and hydrozoan tentacle bulbs as important components of the nervous and sensory systems, being enriched for genes related to nervous system function and sensory processing. Functional similarities (defined by shared GO terms) were greater than expected by chance alone, suggesting that eye-bearing rhopalia and tentacle bulbs are enriched for similar molecular functions including light sensing. Nevertheless, gene expression is largely divergent between species and samples do not clearly cluster by tissue type. This is perhaps not surprising given the enormous evolutionary distances between crown Scyphozoa, Cubozoa, and Hydrozoa, which diverged by at least 505 Mya (Cartwright and Collins, 2007). Rhopalia of *A. aurita* and *T. cystophora* appear to be more similar in gene expression to one another than to *S. tubulosa* tentacle bulbs, consistent with the evolutionary relationships of these species and the likely homology of rhopalia themselves (Marques and Collins, 2005). These similarities could be driven by genes with roles in the cubozoan/scyphozoan nervous system or non-visual sensory structures, such as sensory ciliated epithelia (Nielsen et al., 2019; Parkefelt and Ekström, 2009). However, we did observe numerous examples of convergent gene expression between rhopalia and tentacle bulbs; since rhopalia and tentacle bulbs are not homologous tissues, these genes represent genuine cases of convergence in eye-bearing structures rather than shared ancestry or ontogeny.

### Convergent gene expression identifies candidate genes with roles in phototransduction and eye development

Although the major signature of gene expression in our study was species-specific, we identified numerous homologous genes expressed in eye-bearing tissues across three distantly-related jellyfish lineages, many of which have putative functions related to sensory processing, nervous system development, and cell signalling. Several of these genes are intriguing candidates for roles in medusozoan photoreception and eye development because they are homologs of known bilaterian vision-related genes, including opsins, transcription factors, and lens components.

Overlapping gene sets expressed in convergently-evolved tissues could arise due to chance alone and need not invoke any particular evolutionary mechanism (e.g., Foster et al., 2022).

Consistent with this, we found that overlaps of eye-tissue-upregulated orthogroups were not greater than expected by chance given the number of shared orthogroups between the three species. However, our analysis probably underestimates similarities among species due to low sample sizes (particularly for *T. cystophora*). Additionally, many convergent genes in this study are homologous to vision-related genes from vertebrates and insects, suggesting that evolution repeatedly favors the use of these genes in eyes. These genes could represent convergence at a molecular level, in which members of the same gene family independently gained functions related to visual systems. Examples of this type of convergence could be certain transcription factors or lens-related genes with roles in eye morphogenesis and development.

Alternatively, genes may belong to an ancestral toolkit that forms a component of eyes—a pattern of deep homology. This is supported by our observation that all three jellyfish eye tissues express homologs of canonical phototransduction genes, including opsins, CNG channels, guanylate cyclases, and cGMP-dependent phosphodiesterases. Additionally, there was a strong enrichment of ion channels and calcium-binding genes (e.g., calmodulins) in all three species.

Calcium signalling is a fundamental feature of bilaterian phototransduction (Koch and Dell’Orco, 2013; Nakatani et al., 2002), and this also appears to be the case in cnidarians. These results suggest that ancient phototransduction genes with conserved light-sensing functions have been repeatedly recruited into eyes (Picciani et al., 2021).

### Opsins

The expression patterns and phylogenetic relationships of opsin proteins support a pattern of lineage-specific recruitment of visual opsins, with one possible exception in Scyphozoa and Cubozoa. We found that most *A. aurita* opsins were not closely related to box jellyfish opsins, indicating that scyphozoan and cubozoan eyes use largely different opsin complements. However, one sequence (“scaffold393.g1”) was orthologous to the known lens-eye opsin of *T. cystophora*. This gene was sister to an opsin sequence from the eyeless scyphozoan *Rhopilema esculentum*, and these scyphozoan genes are orthologs of the eye-specific opsins from box jellyfish (Fig. 3A). If this gene has a role related to the eye-spots of *A. aurita*, it may represent a case of parallel recruitment of an opsin ortholog into independently-evolved eyes. An alternative scenario is that this opsin is expressed in the rhopalia but not specifically in the eyes (in which case it may also be expressed in rhopalia of eyeless scyphozoans). The opsin trees from Picciani et al. (2018) and the current study (Fig. 3A) also contain *A. aurita* sequences from other published assemblies, whose expression we could not quantify here. The use of more complete transcriptome and genome assemblies may clarify the full opsin complement of *A. aurita*. At the same time, *A. aurita* is actually a cryptic species complex (Dawson and Martin, 2001; Scorrano et al., 2017), so sequences from different assemblies may belong to different species.

Overall, we find that *S. tubulosa* and hydrozoans in general have a greater diversity of opsins than scyphozoans and cubozoans, and that opsins largely diversified separately in Hydrozoa and Acraspeda (Scyphozoa+Cubozoa+Staurozoa). Nearly all examined opsins in the scyphozoan *A. aurita* and the cubozoan *T. cystophora* had strong rhopalia-specific expression. In contrast, the hydrozoan *S. tubulosa* possesses a higher number of opsins, some of which were expressed specifically in manubria, tentacles, tentacle bulbs, or in both tentacles and tentacle bulbs. This shows that opsins may have diverse tissue-specific functions in hydrozoans, which may include light-mediated processes such as gonad release (Quiroga Artigas et al., 2018), modulation of cnidocyte firing (Picciani et al., 2021; Plachetzki et al., 2012), or functions unrelated to light perception.

Notably, the four *S. tubulosa* opsins with strong tentacle bulb-specific expression are orthologs of *Cladonema radiatum* visual opsins, suggesting that the common ancestor of these species already possessed eyes expressing 2-3 opsins. The *S. tubulosa* opsin with the highest expression in tentacle bulbs is an ortholog of the *C. radiatum* opsin CropJ, which is localized near the retina margin in that species (Suga et al., 2008). Overall, similarities in expression patterns of opsin orthologs provide functional evidence for the homology of eyes in these species and insights into the possible opsin complement of their ancestor, although their expression has yet to be localized specifically to eyes in *S. tubulosa*. Further work to sequence and quantify expression of opsins in other related species whose eyes are not definitively homologous (e.g., *Moerisia spp.* and *Sphaerocoryne spp.*) could clarify the number of eye origins in this group.

### Other eye-related genes

Genes encoding lens proteins were upregulated in *T. cystophora* and *S. tubulosa*. While the former species has true lenses, ocelli of *S. tubulosa* instead contain an electron dense material that may simply provide structure rather than acting as a lens (Singla and Weber, 1982). We found that *T. cystophora* rhopalia upregulated two previously-characterized crystallin genes (J1A and J1B; Piatigorsky et al., 1993, 1989), as well as an *alpha(B)-crystallin*-like heat-shock protein.

Crystallins are a functional class of proteins involved in lens formation and light focusing, and different eye origins usually recruit different crystallins (Oakley, 2024). Crystallins are not known from other cnidarians, and rather than crystallin genes, we retrieved two lens fiber membrane intrinsic protein-like (Lmip-like) genes upregulated in *S. tubulosa* tentacle bulbs. Interestingly, one of the Lmip-like genes was also upregulated in *T. cystophora* rhopalia. Lmip genes encode structural constituents of mammalian eye lenses (Alcala et al., 1975), making these *S. tubulosa* and *T. cystophora* genes intriguing candidates for the convergent recruitment of homologous genes as structural components of eyes. Finally, aquaporins were expressed in rhopalia of *A. aurita* and *T. cystophora*. Aquaporins occur in all animals and are essential components of vertebrate ocular tissues (Schey et al., 2014), again making this gene family a candidate for convergent recruitment into separate eye origins.

We also identified transcription factors related to genes with known roles in bilaterian eye development: six3/6 (optix) homeobox genes and NR2E-like nuclear receptors expressed in all three species, Lhx1 homeobox genes expressed in *A. aurita* and *S. tubulosa*, and a retinal homeobox gene (Rax) expressed in *T. cystophora*. Six3/6 genes regulate eye development in flies and vertebrates (Anderson et al., 2012) and six1/2 and six3/6 are also expressed in *C. radiatum* eyes (Stierwald et al., 2004). Nuclear receptor subfamily 2 group E genes (NR2E) are essential for vision in other animals (Forrest and Swaroop 2012; Kitambi and Hauptmann 2007; Yu et al. 2000), and Lhx1 regulates retinal development and circadian neurogenesis in vertebrates (Bedont et al., 2014; Kawaue et al., 2012). Finally, Rax is another homeobox gene involved in vertebrate eye formation (Rodgers et al., 2018).

Box jellyfish may use eye-specific neuropeptides as neurotransmitters, and some neuropeptides are expressed in specific neuronal subpopulations in *T. cystophora* rhopalia (Nielsen et al., 2021). However, we found that across all three species, neuropeptides were highly expressed in all tissues, suggesting a more general role in nervous system function. This is consistent with prior studies showing RFamide-expressing neurons throughout the medusa—although other neuropeptides may have tissue-specific expression (Nielsen et al., 2021). In *A. aurita* and *T. cystophora*, all investigated neuropeptide genes had the highest expression in rhopalia but were also highly expressed in manubria. This likely reflects the density of neurons in rhopalia rather than a specific role in visual processing. In contrast, the tentacle bulbs of *S. tubulosa* showed less of an enrichment for neuropeptides. Some *S. tubulosa* neuropeptides were most highly expressed in manubria or tentacles rather than tentacle bulbs, again suggesting functions beyond vision.

## 5. Conclusions

Many genes upregulated in eye-bearing tissues in jellyfish are homologs of bilaterian gene families involved in vision, suggesting that cnidarian and bilaterian eyes draw from a shared genetic toolkit to some extent. This likely reflects the deep homology of animal phototransduction systems, as we observed genes related to opsin-CNG phototransduction (e.g., opsins, ion channels, calcium-binding proteins) upregulated in the eye tissues of multiple jellyfish species. Further characterization of ocular and non-ocular phototransduction in Medusozoa is needed to determine which genes had roles in light perception that pre-date eyes, and which have specifically been co-opted into eyes themselves. We also identify several key genes that may affect eye function and development. For instance, six3/6 family transcription factors were upregulated in the eye-bearing tissues of all three species; these genes are expressed in the eyes of another hydrozoan (*C. radiatum*) and are also involved in eye development in *D. melanogaster* and vertebrates, making this gene family an intriguing candidate for functional testing. However, convergently-evolved eyes in different jellyfish classes (Scyphozoa, Cubozoa, and Hydrozoa) were largely distinct transcriptomically, indicating that most eye-related gene expression is lineage-specific. Some level of convergence at the gene level is expected by chance due to the finite size of genomes, which may explain some of the overlapping gene sets expressed in these species. On the other hand, genes that are repeatedly expressed in vertebrate, arthropod, and cnidarian eyes may be influenced by functional constraints or evolutionary dynamics that favor their use in eyes. Eye evolution seems to involve many possible genetic solutions, but draws on some key phototransduction genes and transcription factors that are repeatedly recruited across independent eye origins. As evolutionary biologists gain more empirical knowledge about gene re-use in convergent systems, they will better understand how factors like divergence time and genetic constraints impact the repeatability of evolution.

## Supporting information

Cnidops tree file

## Acknowledgements

We thank all members of the Oakley Lab for valuable discussions and feedback on the manuscript. We especially thank Carolina Camargo, Jessica Goodheart and Lisa Mesrop for advice on RNA extraction and analysis of gene expression. We gratefully acknowledge the technical support of the GeneCore at the European Molecular Biology Laboratory for RNA sequencing. This research was funded by National Science Foundation (NSF) DEB-2153773 awarded to THO. We analyzed data using High Performance Computing resources purchased with funds from the NSF (CNS-1725797) and administered by the Center for Scientific Computing (CSC) at UC Santa Barbara. The CSC is supported by the California NanoSystems Institute and the Materials Research Science and Engineering Center (MRSEC; NSF DMR 2308708) at UC Santa Barbara.This work was supported by CAPES (Coordenação de Aperfeiçoamento de Pessoal de Nível Superior) through a doctoral scholarship to NP (Process BEX-13130-13/7) in the program Science without Borders.

## 6. Authors contributions

NP: Conceptualization, Data Collection and Curation, Formal Analysis, Investigation, Software, Writing – Original Draft Preparation

CAB: Formal Analysis, Investigation, Visualization, Software, Writing – Review & Editing

SN: Formal Analysis, Investigation,Writing – Original Draft Preparation

JM: Data Collection

APO: Data Collection

MS: Data Collection

DA: Data Collection

AG: Data Collection, Writing – Review & Editing

THO: Conceptualization, Funding Acquisition, Supervision, Writing – Review & Editing

## 8. Data availability

Raw RNA-seq data have been uploaded to the NCBI Sequence Read Archive (SRA), Bioproject PRJNA1236395. Data files and code are available on Github at

https://github.com/npicciani/eye-expression.

